# Platelet mitochondrial transfer via extracellular vesicles modulates neutrophil phenotype and function

**DOI:** 10.1101/2025.01.29.635414

**Authors:** Harriet E. Allan, Nicola Dark, Paul Vulliamy, Marilena Crescente, Paul C. Armstrong, Plinio Ferreira, Timothy D. Warner

**Affiliations:** Centre for Immunobiology, Blizard Institute, Queen Mary University of London, Faculty of Medicine and Dentistry, London; Centre for Trauma Sciences, Blizard Institute, Queen Mary University of London, Faculty of Medicine and Dentistry, London; Imperial College London, London, United Kingdom

**Keywords:** Extracellular vesicles, mitochondria, mitochondrial transfer, neutrophils, NETs

## Abstract

Platelet activation causes the release of extracellular vesicles, of which a small proportion contain respiratory competent mitochondria. Mitochondria are integral for energy production and in the regulation of apoptotic pathways. However, the existence of extracellular mitochondria highlights a potential new role in intercellular communication. We hypothesised that platelet extracellular vesicles could be taken up by circulatory cells and alter their function. In this work we demonstrate that platelet extracellular vesicles containing mitochondria interact with and are internalised by neutrophils. Flow cytometry revealed that this interaction promotes neutrophil surface receptor changes, indicative of enhanced neutrophil activation, adhesion and migration pathways. The internalisation of platelet mitochondria renders neutrophils unable to subsequently engulf bacteria, demonstrating reduced phagocytic capacity, but enhances the formation of neutrophil extracellular traps, both alone and in the presence of additional stimuli. Our findings show that platelet mitochondria released in extracellular vesicles can alter neutrophil activity and so may be important intercellular communicators and modulators of inflammatory and immune responses.

## Introduction

Small anucleate platelets are fundamental for physiological haemostasis but also contribute to pathological thrombosis, immunity and infection. Common to platelet activation during both processes is the production of platelet extracellular vesicles (EVs), which are shed from the plasma membrane and released into the circulation.^1–3^ Platelet-derived EVs are normally present in the healthy circulation, demonstrating an important role in physiological intercellular communication through the transfer of bioactive cargo from platelets to circulatory cells.^1,4,5^ It is also well documented that the circulating levels of platelet-derived EVs increase in diseases associated with thrombosis and inflammation.^1,6–9^ Interestingly, much like interactions between platelets and circulating cells, platelet EVs readily interact with leukocytes in pathological conditions, suggesting they may be important in intercellular communication in these scenarios.^10–13^

Consistent with the heterogeneous nature of platelets, platelet EVs are diverse in terms of their receptor repertoire and intracellular contents such as ribonucleic acids, lipids, cytokines, and organelles^14^. Of particular interest to us, recent work has highlighted that platelets are able to release their mitochondria either packaged within EVs or as free organelles.^15–19^ However, the mechanisms underlying platelet mitochondrial release and the consequences of these processes on other circulating cells remain to be fully explored.

Mitochondria are crucial to the maintenance of cellular health, regulating metabolism as well as apoptosis.^20,21^ Recent research has revealed the phenomenon of mitochondrial transfer between cells both *in vivo* and *in vitro*, highlighting a mechanism in which cellular function may be augmented or rescued through metabolic reprogramming and improving respiratory capacity.^22–26^ These transfer processes have garnered much interest and have been described in numerous contexts and cell types including fibroblasts, mesenchymal stem cells, neurons, cardiomyocytes, platelets and tumour cells.^22,26–30^ Platelet mitochondrial transfer has been reported in both wound healing and following myocardial infarction as promoting repair and recovery.^23,31^ On the other hand, mitochondrial transfer from platelets in the context of both lung and breast cancer promotes metabolic reprogramming in tumour cells, enhancing malignancy and promoting metastasis.^27,30^ Notably, upon release from platelets, both free- and extracellular vesicle-encapsulated mitochondria maintain their capacities to respire.^32^ Therefore, if incorporated into vascular or circulatory cells they could augment metabolic function by increasing mitochondrial mass and so alter functionality.^32^ With recent literature pointing to platelet EVs as a ready vehicle for mitochondrial transfer between cells, we were led to question what effect mitochondria-containing platelet vesicles could have on other circulatory cells. Literature suggests that interactions between platelet EVs and circulatory cells are predominantly with leukocytes, however the consequences of these interactions are unclear.^12,13^ In this study, we explored the functional significance of the interactions between neutrophils and platelet EVs containing and not containing mitochondria.

## Materials and Methods

### Ethical statement: Human studies

All studies were conducted according to the principles of the Declaration of Helsinki and approved by St Thomas’s Hospital Research Ethics Committee (Ref. 07/Q0702/24) or Queen Mary University of London Research Ethics Committee (Ref. QME24.0294). All volunteers gave written informed consent.

### Blood collection and isolation of platelets and neutrophils

Blood was collected by venepuncture into tri-sodium citrate (3.2%; Sigma) from healthy volunteers (aged 25–48), and platelet rich plasma (PRP) obtained by centrifugation at 175 *xg*, for 15 min, with no brake.

### Washed platelet isolation

Platelets were further purified as previously described.^33^ Briefly, washed platelets were prepared by centrifugation of PRP (1000 *xg*, 10 min) in the presence of prostacyclin (PGI_2_, 2μmol/L; Tocris) and apyrase (0.02U/mL, Sigma) and resuspension in modified Tyrode’s HEPES (MTH) buffer (134mmol/L NaCl, 2.9mM KCl, 0.34mmol/L Na_2_HPO_4_, 12mmol/L NaHCO_3_, 20mmol/L HEPES and 1mmol/L MgCl_2_; pH 7.4; Sigma), with glucose (0.1% (w/v); Sigma). Washed platelets were adjusted to a concentration of 3 × 10^8^/ml, rested for 30min and were supplemented with calcium chloride (CaCl_2_, 2mmol/L; Sigma) prior to use.

### Neutrophil isolation

Following the removal of the PRP layer, red blood cells were sedimented using dextran (6%, Thermo Fisher Scientific) and the leukocytes were separated using centrifugation (400 xg, 30 min) over Histopaque 1077 (Sigma) and lysis with ice cold dH_2_O. The sample was supplemented with Hank’s Balanced Salt Solution (Sigma) and phosphate buffered saline without calcium or magnesium (PBS^-/-^, Sigma) washed by centrifugation (300 *xg*, 10 min) and resuspended in RPMI without phenol red (Sigma).

### Immunofluorescence and confocal microscopy of activated platelet samples

Washed platelets (3 × 10^8^/ml) were stained with MitoTracker Orange CMTMRos (20nm; Life Technologies) for 15min and supplemented with eptifibatide (4µM, Sigma) for 5 min. Samples were incubated with vehicle or thrombin receptor activating peptide SFLLRN (TRAP-6 amide, 20µM; Bachem) in a Born Light Transmission Aggregometer (5min, 1200rpm, 37°C). Samples were fixed with paraformaldehyde (PFA, 4%; VWR) for 10 min and prepared for immunofluorescence as previously described.^34^ Briefly, fixed platelets were centrifuged onto poly-l-lysine coated coverslips (VWR; 600 *xg*, 5 min), permeabilized with 0.2% Triton, 2% donkey serum, 1% bovine serum albumin (BSA) in PBS (all Sigma) and incubated with mouse monoclonal anti-α-tubulin (1:200; Sigma) and subsequently anti-mouse Alexa Fluor 488 (1:500, Thermo Fisher Scientific). Samples were mounted with Prolong Diamond antifade mount (Thermo Fisher Scientific). Confocal microscopy was performed using an inverted Zeiss LSM880 with Airyscan confocal microscope; 63x objective, 1.4 Oil DICII (Zeiss). Analysis was conducted using Zen Software (2.3 SP1, Zeiss) and ImageJ (NIH).

### Production and characterisation of platelet extracellular vesicles

Platelet EVs were produced as previously described.^35^ Briefly, washed platelets (3 × 10^8^/ml) were stained with MitoTracker Orange CMTMRos (20nm) for 15min and statically incubated with the synthetic peptide TRAP-6 amide (20µM) for 2 hours at 37°C. Residual platelets were removed by double centrifugation (1000 *xg*, 10min), the supernatant collected and the EVs were pelleted at 25,000 *xg*, 45min. The platelet EV pellet was stored at -80°C until required.

### Surface receptor expression

Platelet EVs were characterised by imaging flow cytometry (Image Stream^x^ MKII) as previously described.^36^ Briefly, samples were stained with a CD41 APC (1:200; clone HIP8; Biolegend) and Cell Trace Violet (1:50, Thermo Fisher Scientific) or CD62P BV421 (1:100; clone AK4; Biolegend) or Annexin V Pacific Blue (1:100; Biolegend) for 30 min. Samples were analysed using the Image Stream^x^ MKII collecting 10,000 events. Analysis was performed using IDEAS software and FlowJo v.10.

### Respiratory capacity

The respiratory capacity of platelet EVs, determined by oxygen consumption rate, was assessed using an Agilent Seahorse XF24 (Agilent). The XF24 plate was coated with CellTak (2.4µg/m) for 20 min and the cartridge was prepared as detailed in the manufacturer’s instructions. Platelet EVs were prepared as described with the modification of resuspension in warmed Seahorse base medium (2mM glutamine, 25mM glucose, 1mM pyruvate). Platelet EVs (3 × 10^8^) and washed platelets (6 × 10^7^) were spun onto the CellTak coated plate (200 *xg*, 1 min), allowed to rest for 30 min at 37°C and subsequently placed in the XF24 Seahorse Analyser to measure oxygen consumption rate for a further 30 min.

### Characterisation of platelet extracellular vesicles interactions with neutrophils

Platelet EVs stained with MitoTracker Orange CMTMRos (20nm) and BODIPY (2µm, Thermo Fisher Scientific) were incubated with isolated neutrophils (5 × 10^6^/ml) to assess EV-neutrophil and mitoEV-neutrophil interactions. Samples were fixed with PFA (4%) after 5, 30 and 60 min to determine the rate of interactions. Following fixation, samples were spun onto poly-l-lysine coated coverslips (600 *xg*, 5 min), blocked and permeabilised with 0.2% Triton, 2% donkey serum, 1% BSA in PBS, and stained with Phalloidin Alexa Fluor 647 (1:200; Thermo Fisher Scientific) and DAPI (1:10,000, Abcam). Confocal microscopy was performed using an inverted Zeiss LSM880 with Airyscan confocal microscope; 63x objective, 1.4 Oil DICII (Zeiss). Analysis was conducted using Zen Software (2.3 SP1, Zeiss) and ImageJ (NIH).

Neutrophil mitochondrial mass and function was assessed following incubation with platelet EVs, using MitoTracker Green FM (50nm, mitochondrial mass; Thermo Fisher Scientific) and MitoTracker Deep Red (50nm, functional mitochondria; Thermo Fisher Scientific). Samples were analysed by flow cytometry (BD LSRII) and analysis was performed using BD FACs Diva and FlowJo v.10.

### Isolation of platelet extracellular vesicle subpopulations

Cell sorting (BD FACs Aria III Fusion Cell Sorter; 70µm nozzle, 70Ps) was used to separate mitochondria containing platelet EVs (mitoEV; MitoTracker Orange CMTMRos positive; ∼20% of the total EV population) from the rest of the extracellular vesicle population (EV; MitoTracker Orange CMTMRos negative; Supplemental Figure 1). Samples were subsequently pelleted at 25,000 *xg*, 45 min and stored at -80°C until required.

### Assessing changes in neutrophil surface receptor expression

Isolated neutrophils (5 × 10^6^/ml) were incubated alone, or with EVs or mitoEVs for 60 min at 37°C and subsequently analysed by flow cytometry to determine changes in receptor expression. Samples were stained with antibodies against CD66b APC (1:200; clone G10F5), CD11b Pacific Blue (1:200; clone ICRF44), CD192 Brilliant Violet 421 (1:100; clone K036C2), CD62L Pacific Blue (1:100; clone DREG-56), CD31 AF647 (1:100; clone WM59), CD16 Pacific Blue (1:100; 3G8), CXCR2 Pacific Blue (1:100; clone 5E8/CXCR2), CD36 APC (1:100; clone 5-271), ICAM-1 Pacific Blue (1:100; clone HCD54) or Annexin V APC (1:50) (all Biolegend), and 10,000 events were acquired on flow cytometer (BD LSRII). For a subset of markers, mitoEVs were pre-treated with carbonyl cyanide-p-trifluoromethoxyphenylhydrazone (FCCP; 1µM, Sigma) for 30min, washed by centrifugation (25,000 xg, 45min) prior to incubation with isolated neutrophils. Analysis was performed using BD FACs Diva and FlowJo v.10. Analysis was performed using BD FACs Diva and FlowJo v.10.

### Determining changes in neutrophil intracellular calcium and reactive species generation

Isolated neutrophils (5 × 10^6^/ml) were incubated alone, with EVs or mitoEVs for 60 min at 37°C and subsequently stained with CD66b Pacific Blue (1:100; clone G10F5; Biolegend) and Calbryte^630^ (5µm, Stratech) or CellRox Deep Red (500nm; Thermo Fisher Scientific). Samples acquisition was performed by flow cytometry (BD LSRII) collecting 10,000 CD66b-positive events. Analysis was performed using BD FACs Diva and FlowJo v.10.

### Characterisation of changes in neutrophil function

#### Neutrophil extracellular Trap formation

Neutrophil extracellular Trap (NET) formation was assessed under basal conditions, as well as in the presence of lipopolysaccharide (LPS, 25ng/ml; Sigma) or phorbol 12-myristate 13-acetate (PMA, 200nM; Sigma). Briefly, neutrophils were incubated alone, with EVs or mitoEVs in presence of vehicle (PBS), LPS or PMA in poly-l-lysine coated ibidi chambers for 90 min at 37°C. Samples were subsequently fixed with PFA (4%), blocked and permeabilised and stained with anti-rabbit myeloperoxidase (MPO, 1:200; Agilent), followed by anti-rabbit Alexa Fluor 647 (1:500; Thermo Fisher Scientific) and DAPI (1:10,000). NET formation was assessed by immunofluorescence using a Zeiss LSM880 confocal microscope; 20x objective, 1.4 DICII (Zeiss). Analysis of DNA-length in the NETs was conducted using Zen Software (2.3 SP1, Zeiss) and ImageJ (NIH).

#### Phagocytosis

Neutrophil phagocytic ability was assessed using pHrodo™ *E*.*coli* Bioparticles and flow cytometry (BD LSRII). Briefly, neutrophils were incubated with EVs or mitoEVs for 60 min, followed by pHrodo™ *E*.*coli* Bioparticles (Thermo Fisher Scientific) according to manufacturer’s guidelines and CD66b Pacific Blue (1:100; Biolegend). Using flow cytometry (BD LSRII), 10,000 CD66b positive events were captured and the degree of phagocytosis was determined by fluorescence of the pHrodo™ *E*.*coli* Bioparticles.

### Statistical Methods

Data are expressed as mean ± SEM. Graphs and statistical analysis were generated using GraphPad Prism 9 (GraphPad Software Inc.). Statistical analyses were performed with a paired t-test or a one-way ANOVA with Dunnett’s post-test for multiple comparisons as appropriate. Significance was defined as p < 0.05.

## Results

### Platelet activation causes release of mitochondria within extracellular vesicles

Incubation of platelets with TRAP-6 amide caused a rapid and significant reduction in the number of mitochondria within platelets, such that after 5 min the number had fallen from 7±1 mitochondria per platelet to 4±0.5 (n=6, p<0.01; Figure 1A-C). This reduction was accompanied by a significant decrease in platelet area from 6.0±0.4µm^2^ to 4.3±0.2µm^2^ per platelet (p<0.01; Figure 1D) compatible with the formation of a heterogenous population of EVs. Imaging flow cytometry demonstrated CD41+ platelet-derived vesicles with high levels of Annexin V binding (82±2%) and moderate levels of P-selectin expression (30±4%). Mitochondria were encapsulated within 21±2% of these platelet-derived EVs (Figure 2A-B). Mitochondria-containing EVs (mitoEVs) were of larger size and 95±1% positive for P-selectin (Figure 2C; Supplemental Figure 2). Mitochondria within these EVs consumed oxygen at a rate of 36±7.5pmol/min (per 3 × 10^8^ EVs; n=5) compared to 74±8pmol/min in intact washed platelets (per 6 × 10^7^ platelets; n=5; Figure 2D).

**Figure 1:**
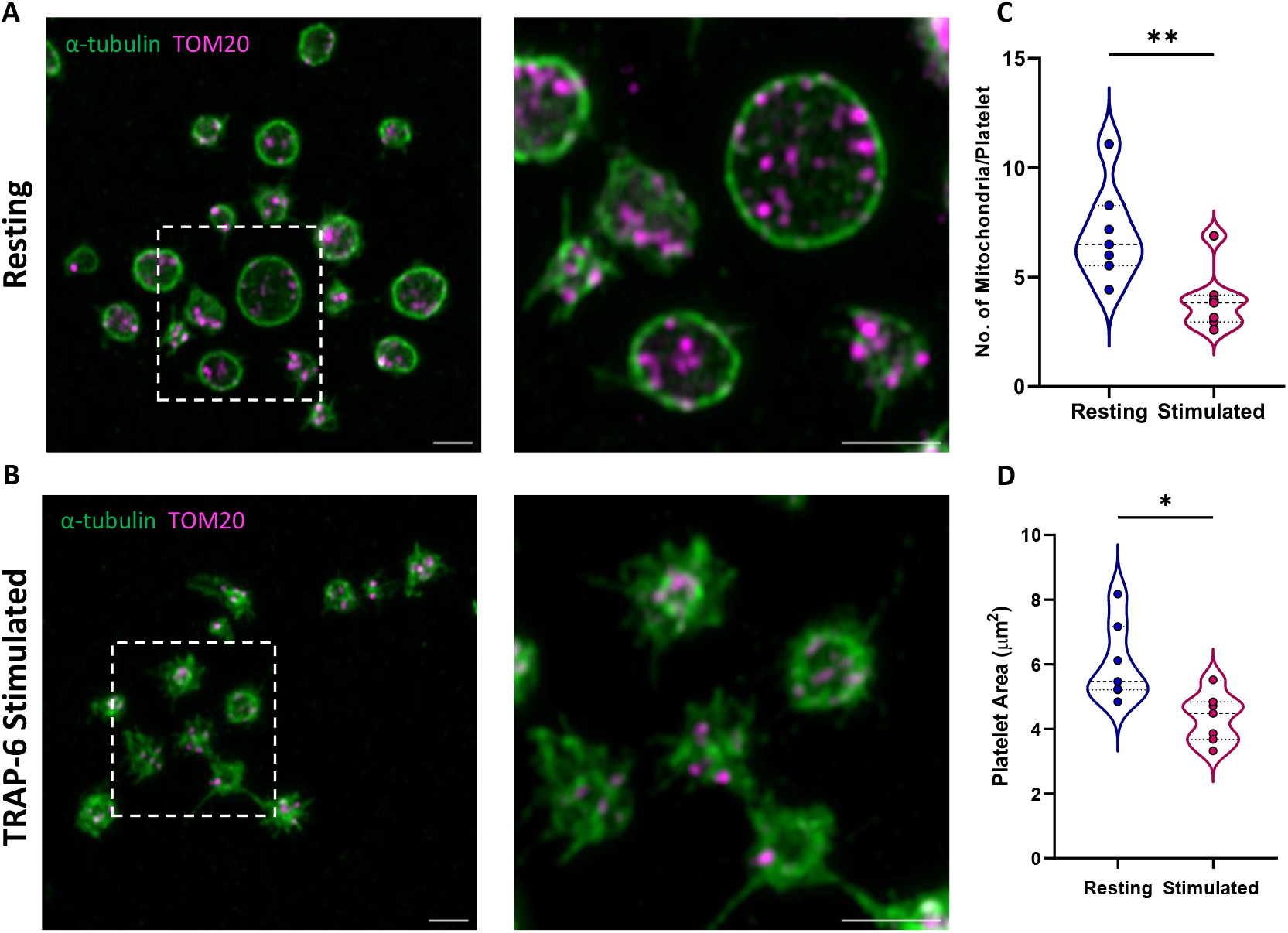
Characterisation of platelet mitochondria following activation. Representative confocal images showing platelets incubated with **(A)** vehicle and **(B)** TRAP-6 stained with α-tubulin (green) and TOM20 (magenta). White insert boxes show a zoomed-area at 10x magnification. Scale bars represent 2µm. Quantification of **(C)** mitochondrial number and **(D)** platelet area in platelets incubated with vehicle (blue) and TRAP-6 (magenta). Data presented as violin plots with individual values and line at median and quartiles, n=6 (*p<0.05, **p<0.01).

**Figure 2:**
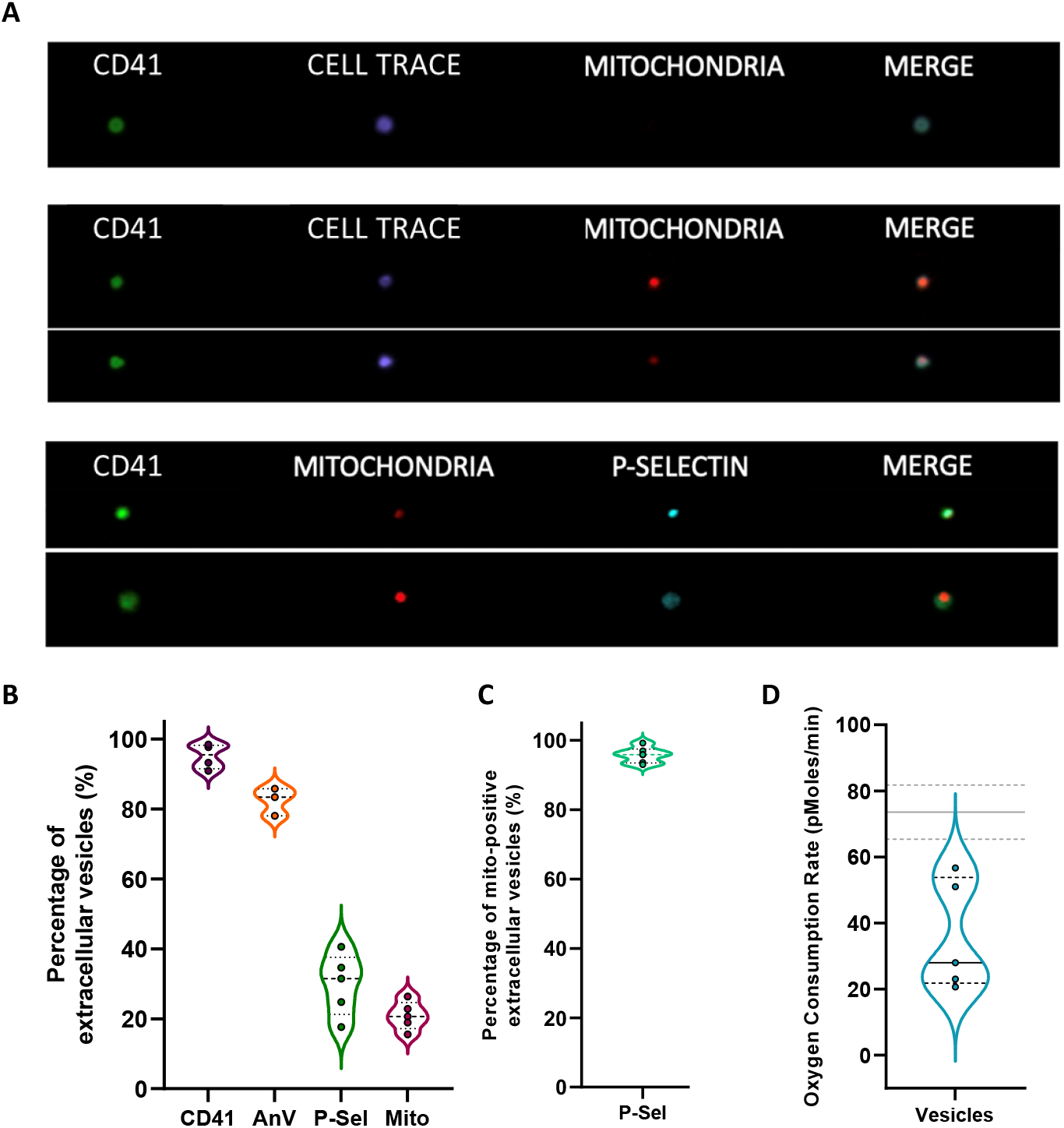
Characterisation of platelet-derived extracellular vesicles following incubation with TRAP-6. **(A)** Representative image stream pictures showing platelet extracellular vesicles stained for CD41 (green), cell trace (purple) to stain the cytoplasm, MitoTracker Orange (red) and P-selectin (cyan). **(B)** Quantification of the percentage of extracellular vesicles positive for CD41 (purple), Annexin V (orange), P-selectin (green) and MitoTracker (magenta). **(C)** Percentage of P-selectin expression on MitoTracker positive extracellular vesicles. **(D)** Oxygen consumption rate (OCR) as an indicator of respiratory capacity in platelet extracellular vesicles, grey lines indicate OCR in platelets. Data presented as violin plots with individual values and line at median and quartiles, n=4.

### Platelet mitochondria EVs bind to and are internalised by neutrophils

Confocal microscopy revealed that mitoEVs bound to neutrophils in a time-dependent manner over 60 minutes (Figure 3A-B). MitoEVs initially bound to the periphery of the cell with internalisation being observed after 30 minutes. Accompanying these interactions was an increase in neutrophil area, from 36±4µm^2^ per neutrophil under basal conditions to 47±4µm^2^ and 59±3µm^2^ at 30 minutes and 60 minutes respectively (p<0.05; n=4; Figure 3C).

**Figure 3:**
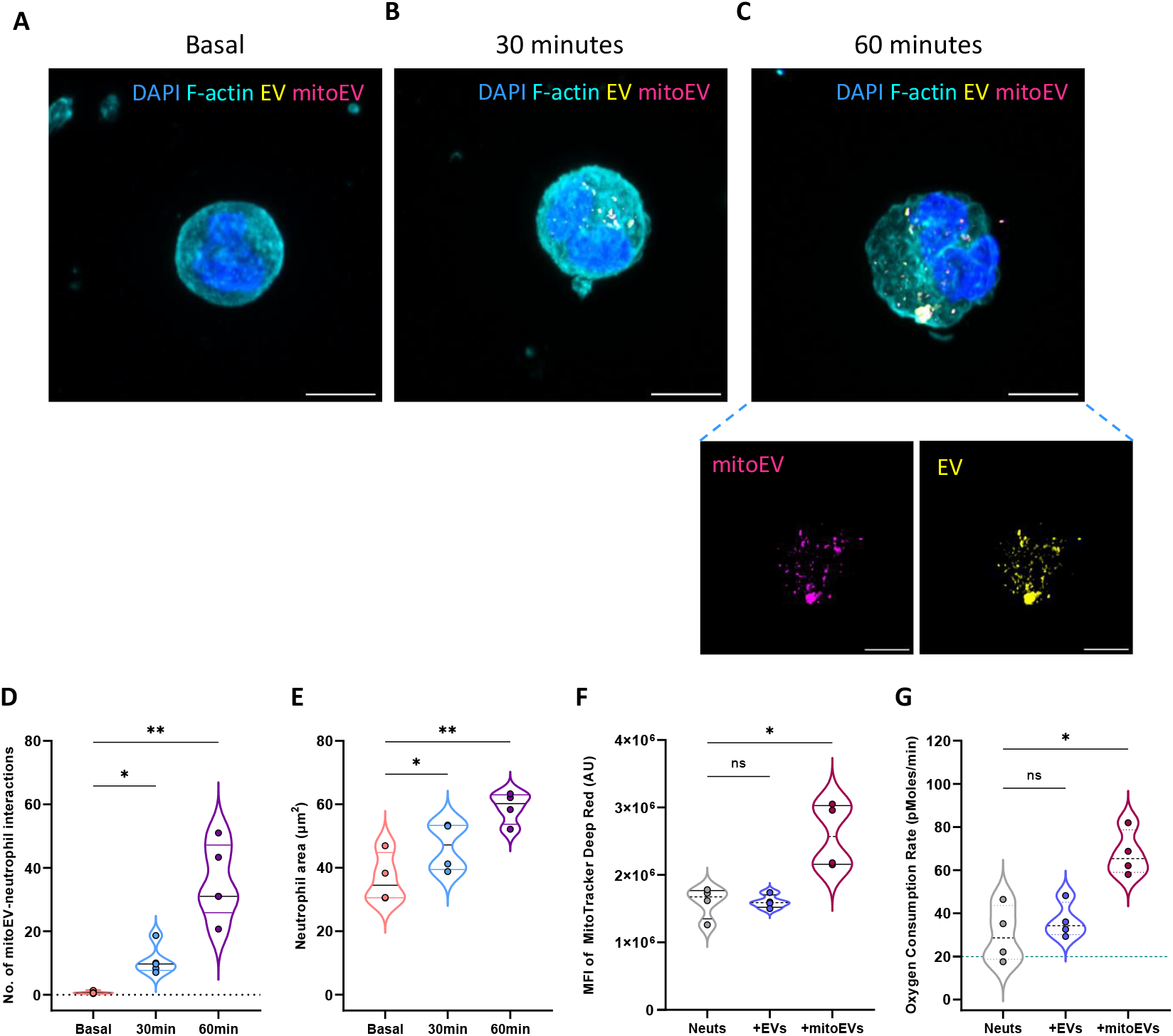
Characterisation of platelet extracellular vesicle interactions with neutrophils. Representative confocal microscopy images showing platelet extracellular vesicles (EVs; yellow) with mitochondria (mitoEV; magenta) interacting with neutrophils stained for F-actin (cyan) and DAPI (blue) **(A)** under basal conditions and after co-incubation for **(B)** 30 and **(C)** 60 minutes (insert indicates separated vesicles (yellow) and mitochondria vesicles (magenta). White represents the colocalization of EVs and mitoEVs. Scale bar represents 5µm. **(D)** Quantification of the number of mitochondria-containing platelet vesicles (mitoEV) interacting with neutrophils at basal, 30 and 60 minutes. **(E)** Quantification of the neutrophil area under basal and following 30 and 60 minutes incubation with platelet vesicles. **(F)** Quantification of neutrophil mitochondrial mass following incubation with platelet vesicles containing a mitochondria (mitoEV) and those without (EV) for 60 minutes. **(G)** Oxygen consumption rate (OCR) of neutrophils alone (grey), and those incubated with EVs (blue) or mitoEVs (magenta) for 60 minutes. Data presented as violin plots with individual values and line at median and quartiles, n=4 (*p<0.05, **p<0.01).

### Internalisation of mitochondria EVs increases oxidative phosphorylation in neutrophils

Following isolation of two subpopulations of platelet EVs; those containing mitochondria (mitoEV) and those without mitochondria (EV) the differential effects on neutrophils were explored. Incubation of neutrophils with mitoEVs but not with EVs lacking mitochondria caused a significant increase in the neutrophils staining for MitoTracker Deep Red (Figure 3D) and MitoTracker Green (Supplemental Figure 3). Similarly, following incubation with mitoEVs, but not EVs, there was a significant increase in neutrophil oxygen consumption rate from 30.3±6.5pmol/min in neutrophils alone to 67.7±.2pmol/min in neutrophils + mitoEVs (p<0.05; n=4; Figure 3E) which did not occur when neutrophils were incubated with EVs

### Binding of platelet mitochondria vesicles causes changes in neutrophil receptor expression, intracellular calcium levels and reactive oxygen species levels

Incubation of neutrophils with mitoEVs but not EVs, caused an increase in neutrophil surface levels of CD66b (126±5%), CD11b (132±8%), CD192 (167±16%) (Figure 4A-C; n=5, p<0.05). These increases were accompanied by reductions in the expressions of CXCR2 (84±3%), CD36 (72±8%) and ICAM-1 (90±2%) (Figure 4D-F; n=5, p<0.05). No changes in neutrophil receptor expression were seen when mitoEVs with pre-incubated with FCCP to cause disruption of the mitochondrial membrane potential (Figure 4G-I).

**Figure 4:**
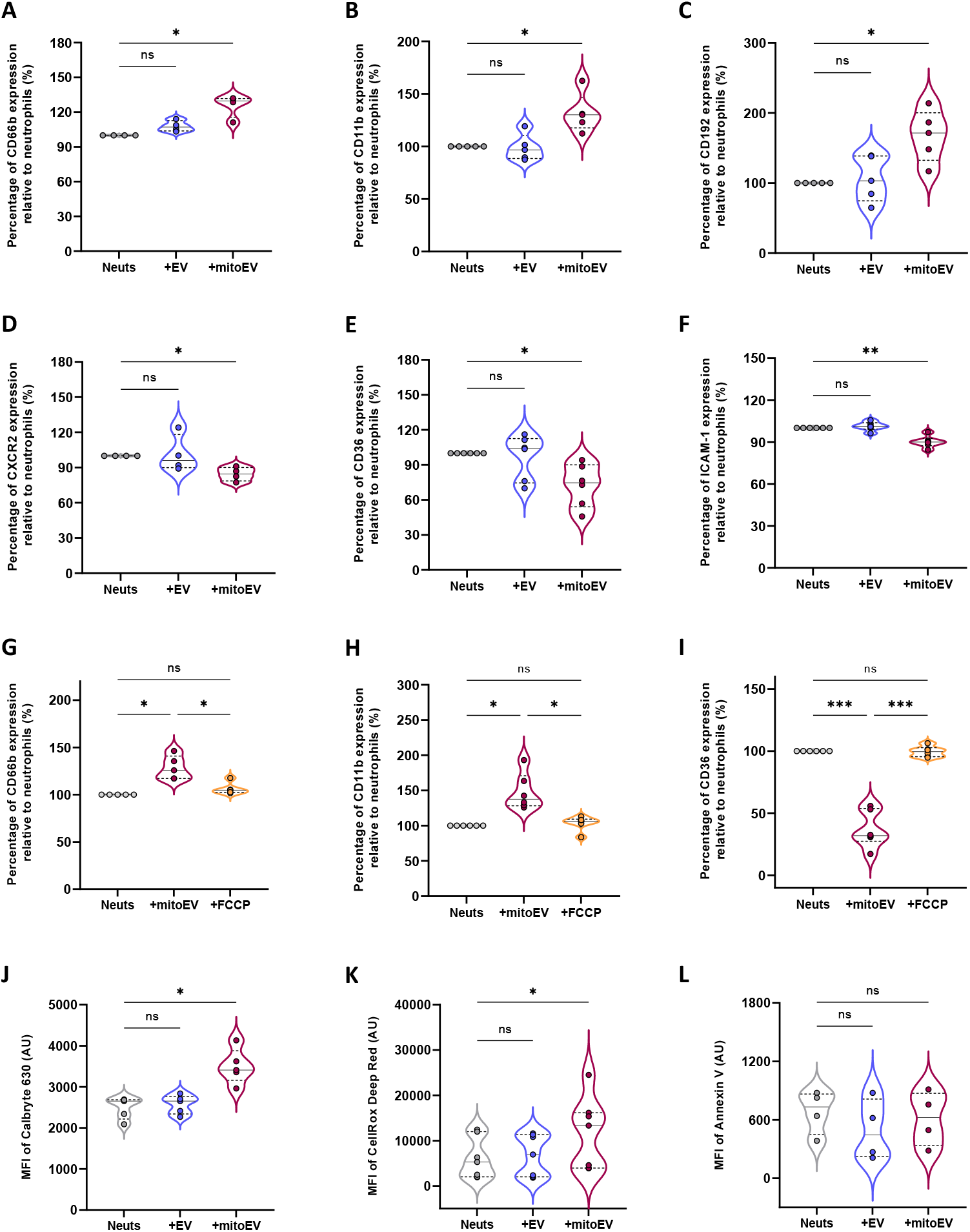
Characterisation of neutrophil surface receptor and intracellular calcium and reactive oxygen species following interactions with platelet extracellular vesicle subpopulations. Flow cytometric analysis of the surface receptor expression of **(A)** CD66b, **(B)** CD11b, **(C)** CD192, **(D)** CXCR2, **(E)** CD36, **(F)** ICAM-1, following incubation for 60min with EVs (blue) or mitoEVs (magenta) relative to neutrophils alone (grey). Analysis of selected surface markers **(G)** CD66b, **(H)** CD11b, **(I)** CCD36 in following incubation of neutrophils with mitoEV (magenta) or mitoEVs treated with FCCP (orange). Flow cytometric analysis of intracellular **(J)** cytoplasmic calcium levels (Calbryte 630), and **(K)** reactive oxygen species (Cell Rox) in neutrophils alone (grey) or following incubation with EVs (blue) or mitoEVs (magenta). **(L)** Analysis of annexin V binding in neutrophils alone (grey) or following incubation with EVs (blue) or mitoEVs (magenta). Data presented as violin plots with individual values and line at median and quartiles, n=6 (*p<0.05, **p<0.01, ***p<0.005).

Incubation of neutrophils with mitoEVs but not EVs caused an increase in cytoplasmic calcium levels (Calbryte^630^ fluorescence; 2490±119AU vs. 3496±191AU in neutrophils vs. neutrophils + mitoEVs; Figure 4J, n=5, p<0.05) and an increase in intracellular reactive oxygen species (CellRox fluorescence; 6060±1711AU vs. 11679±2975AU in neutrophils vs. neutrophils + mitoEVs; Figure 4K, n=5, p<0.05). No differences in annexin V binding were noted between samples (Figure 4L).

### Platelet mitochondria extracellular vesicles reduce neutrophil phagocytic ability but enhance neutrophil extracellular trap (NET) formation

MitoEVs enhanced the capacity of neutrophils to form NETs under basal conditions (neutrophil DNA area from 47±4µm^2^ to 92±14µm^2^; n=4, p<0.05), whilst EVs were without effect (47±5µm^2^, p>0.05, Figure 5A,D). In addition, mitoEVs increased NET formation when combined with either LPS (LPS-stimulated DNA area, 107±21µm^2^ vs. 238±6µm^2^; neutrophils vs. neutrophil + mitoEVs, p<0.05; Figure 5B,E) or PMA (PMA-stimulated DNA area, 137±23µm2 vs. 240±41µm2; neutrophils vs. neutrophils + mitoEVs, p<0.05; Figure 5C,F). Conversely, the interaction with mitoEVs caused a reduction in neutrophil phagocytic capacity, with mitoEV-neutrophils phagocytosing 52±10% of pHrodo™ *E. coli* Bioparticles compared to neutrophils alone (n=6, p<0.05; Supplemental Figure 4).

**Figure 5:**
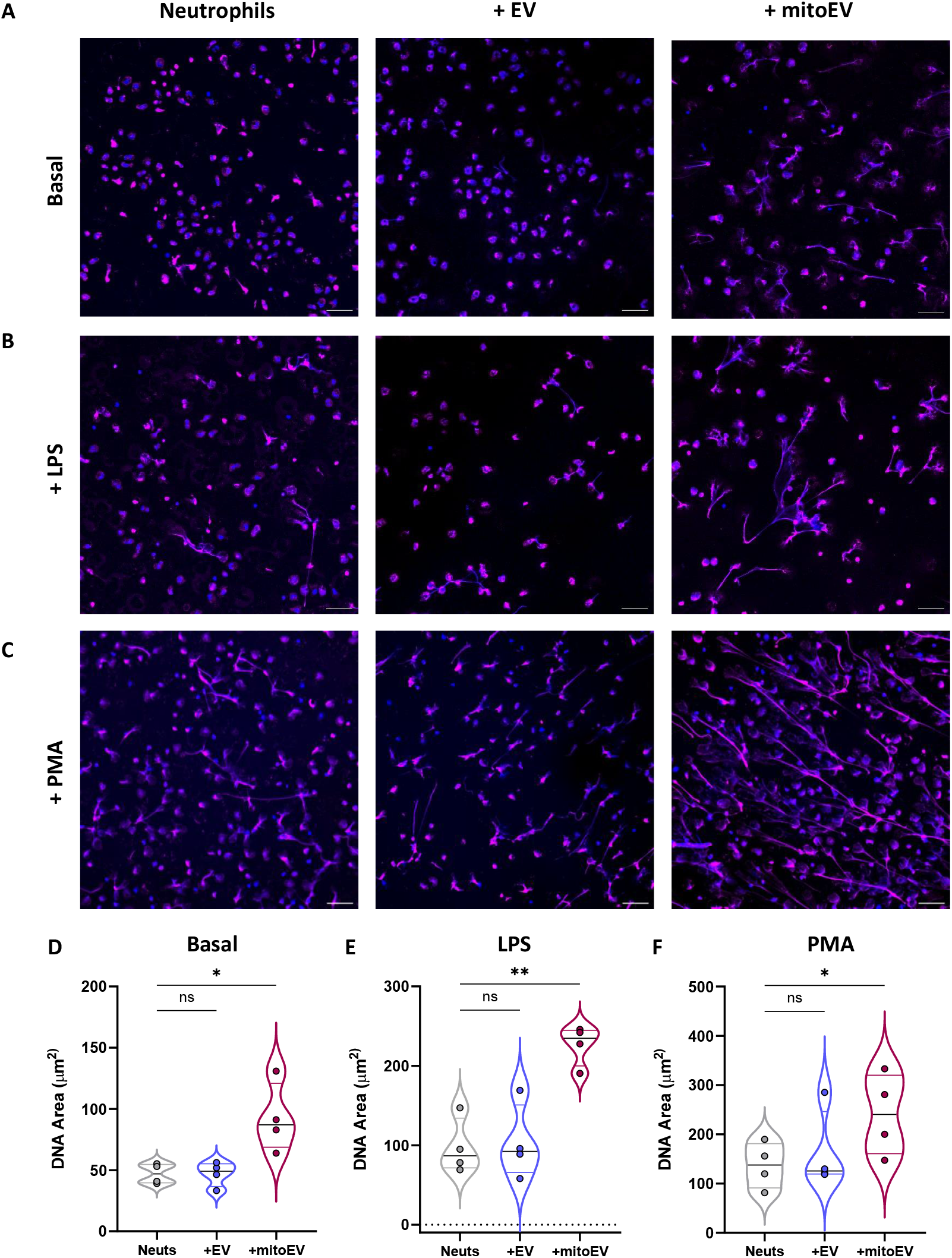
Characterisation of Neutrophil Extracellular Trap formation. Representative confocal microscopy images of neutrophils under **(A)** basal conditions or incubated with **(B)** LPS or **(C)** PMA either alone, +EVs or +mitoEVs, stained with DAPI (blue) and myeloperoxidase (magenta). Scale bar represents 50µm. Quantification of the DNA area under **(D)** basal conditions, **(E)** incubated with LPS and **(F)** incubated with PMA (neutrophils alone, grey; +EVs, blue; +mitoEVs, magenta. Data presented as violin plots with individual values and line at median and quartiles, n=6 (*p<0.05, **p<0.01).

## Discussion

While it has been known for some time that the platelet extracellular vesicle population is heterogeneous, most research has focussed on characterising them as a single population. Here we demonstrate platelet stimulation causes a reduction in the number of intra-platelet mitochondria and an increase in extracellular vesicle production and that mitochondria are present in approximately 20% of the platelet EV population. ^15,16^ These mitochondria maintain their respiratory capacity, highlighting their potential role in intercellular communication. In addition, mitoEVs had higher levels of P-selectin expression than mitochondria-negative vesicles, which led us to speculate that mitoEVs are more likely to interact with and have a functional effect on leukocytes, in particular neutrophils.

Based on these findings, we isolated two subpopulations of platelet EVs, those containing mitochondria and those without, and used them to study the functional effects of mitoEVs and mitochondrial transfer from platelets to neutrophils. Interestingly, we found that mitoEVs, but not EVs enhance neutrophil respiratory capacity, suggesting they may be facilitating metabolic reprogramming and thereby increasing the propensity for inflammatory effector functions. Indeed, mitoEVs efficiently interact with and become internalised by neutrophils and subsequently enhance oxidative phosphorylation suggesting that the mitochondria merge with the existing neutrophil mitochondrial network. Neutrophils are predominantly glycolytic cells, with very low levels of oxidative phosphorylation, and therefore the presence of platelet mitochondria and the additional energy that they confer, may be facilitating metabolic reprogramming and enhancing the neutrophils’ activation pathways.^37^ Interestingly, mitochondrial ATP production has been shown to trigger the first phases of purinergic signalling in neutrophils, initiating their functional responses, which are later supported by ATP production from glycolysis.^38^ In the circumstances presented here, mitochondrial transfer from platelet EVs could prime these processes, facilitating a stronger and more rapid neutrophil activation response.

Neutrophils have a myriad of effector functions, which will be supported by enhanced metabolic function. Indeed, we found that mitoEVs altered the neutrophil receptor repertoire by increasing surface expression of several activation marker including CEACAM8 (CD66b), Mac-1 (CD11b) and chemokine receptor 2 (CCR2; CD192) which are important molecules to promote neutrophil adhesion, rolling and migration suggesting that transfer of platelet mitochondria may be causing neutrophil activation and subsequent migratory behaviour.^39–41^ Supporting this activated phenotype induced by mitoEVs we demonstrate that these neutrophils exhibited higher levels of intracellular calcium which is fundamental for a plethora of neutrophil activation pathways.^42^ One such immune modulatory process which is closely linked with these intracellular increases is the formation and release of NETs.^43–47^ Under basal conditions mitoEVs, but not EVs, promoted the formation of NETs, an effect that was further enhanced in the presence of either LPS or PMA. Supporting the formation of NETs is an increase in reactive oxygen species generation in the presence of mitoEVs which has been shown to be critical for the initiation of NET formation.^48,49^ Emerging evidence suggests that neutrophils are unable to perform both NET formation and phagocytosis, and interestingly, we found that several receptors, including CXCR2, CD36 and ICAM-1, which have been implicated in neutrophil phagocytic behaviour, were reduced following incubation with mitoEVs.^50,51^ Consistent with these receptor changes, we found that neutrophils incubated with mitoEVs, but not EVs, exhibited a reduction in their phagocytic ability. This interesting observation suggests that the internalisation of platelet mitochondria and subsequent enhanced respiratory capacity may be pushing neutrophils to adopt a NET-forming rather than phagocytic phenotype.^52^ The need for transferred mitochondria to be respiratory competent to produce these changes in neutrophils was confirmed by pre-treating the extracellular vesicles with FCCP, which abolishes mitochondrial membrane potential and abolished the effects of mitoEVs on neutrophil receptor expression.

Elevated levels of platelet EVs and extracellular mitochondria have been described in settings of thrombosis, inflammation and tissue injury, as well as in transfusion storage bags, making it important to understand their potential effects on other cells.^1,13,21,32,53–55^ It is clear that the release of mitochondria and mitochondrial transfer is a complex phenomenon and is context and cell type dependent. However, given that they are readily available in transfusion bags, they may present a potential therapeutic strategy for augmenting cellular function in certain scenarios. For instance, transfusion with mitoPMVs in patients with microbial infections, may support the mechanisms involved in NET release thereby promoting the eradication of pathogens from the circulation.^56,57^ Thus, mitochondrial transfer from platelets to neutrophils could help the host to fight infections. However, such methods would need to be used with caution, as increased NET release may have more complex consequences in patients with additional clinical risk factors including thrombosis or vascular disease.^58^ The possible beneficial or detrimental impact associated with increased neutrophil activation and release of NETs consequent to platelet mitochondrial transfer remains to be determined in specific diseases and in the context of transfusion medicine. In addition, the functional effects of mitoEVs on other vascular and circulatory cells would need to be fully explored as previous work has highlighted platelet EVs readily interact with and transfer cargo to monocytes and endothelial cells.^13,59^ However, the efficacy of platelet mitochondrial transfer has recently been demonstrated in acute STEMI patients who have received autologous intracoronary injections of platelet-derived mitochondria and shown favourable outcomes with increased left ventricular ejection fraction and subsequent exercise capacity.^31^

In conclusion, our work demonstrates that platelet mitochondria transfer via EVs enhances neutrophil respiratory capacity and alters their phenotype and function. Our findings support previous research showing that released platelet mitochondria interact with neutrophils and monocytes and offer new insights into the mechanisms by which activated platelets modulate behaviour of these cells and influence immune responses.^15,18^ In addition, our findings further illustrate the heterogeneity among EVs released from activated platelets and demonstrate that this variability in EV cargo is of functional importance for downstream effects on the phenotype and behaviour of recipient cells. Overall, our research builds on previously published work and provides detailed characterisation of the functional and phenotypic changes occurring in neutrophils following platelet mitochondrial transfer.

## Author contributions

HEA designed the research, performed the assays and collected data, analysed and interpreted data, performed statistical analysis, and wrote the manuscript. ND and PF performed the assays, collected data and reviewed the manuscript. PV, MC and PCA, analysed and interpreted data, and revised the manuscript. TDW designed the research, analysed and interpreted data, and revised the manuscript.

## References

1. Melki, I., Tessandier, N., Zufferey, A. & Boilard, E. Platelet microvesicles in health and disease. Platelets 28, 214–221 (2017).

2. Antwi-Baffour, S. et al. Understanding the biosynthesis of platelets-derived extracellular vesicles. Immun Inflamm Dis 3, 133–40 (2015).

3. Yuana, Y., Sturk, A. & Nieuwland, R. Extracellular vesicles in physiological and pathological conditions. Blood Rev 27, 31–39 (2013).

4. Badimon, L., Suades, R., Fuentes, E., Palomo, I. & Padró, T. Role of Platelet-Derived Microvesicles As Crosstalk Mediators in Atherothrombosis and Future Pharmacology Targets: A Link between Inflammation, Atherosclerosis, and Thrombosis. Front Pharmacol 7, 1–17 (2016).

5. Heijnen, H. F. G., Schiel, A. E., Fijnheer, R., Geuze, H. J. & Sixma, J. J. Activated Platelets Release Two Types of Membrane Vesicles: Microvesicles by Surface Shedding and Exosomes Derived From Exocytosis of Multivesicular Bodies and []-Granules. Blood 94, 3791–3799 (1999).

6. Herring, J. M., McMichael, M. A. & Smith, S. A. Microparticles in Health and Disease. J Vet Intern Med 27, 1020–1033 (2013).

7. Puhm, F., Boilard, E. & MacHlus, K. R. Platelet extracellular vesicles; beyond the blood. Arteriosclerosis, Thrombosis, and Vascular Biology vol. 41 87–96 (2021).

8. Zaldivia, M. T. K., McFadyen, J. D., Lim, B., Wang, X. & Peter, K. Platelet-Derived Microvesicles in Cardiovascular Diseases. Front Cardiovasc Med 4, 74 (2017).

9. Vajen, T., Mause, S. F. & Koenen, R. R. Microvesicles from platelets: novel drivers of vascular inflammation. Thromb Haemost 114, 228–36 (2015).

10. Weiss, R. et al. Differential Interaction of Platelet-Derived Extracellular Vesicles with Leukocyte Subsets in Human Whole Blood. Sci Rep 8, (2018).

11. Forlow, S. B., McEver, R. P. & Nollert, M. U. Leukocyte-Leukocyte Interactions Mediated by Platelet Microparticles under Flow. Blood; 95(4): 1317–1323 (2000).

12. Chimen, M. et al. Appropriation of GPIbα from platelet-derived extracellular vesicles supports monocyte recruitment in systemic inflammation. Haematologica 105, 1248–1261 (2020).

13. Vulliamy, P. et al. Histone H4 induces platelet ballooning and microparticle release during trauma hemorrhage. Proc Natl Acad Sci USA 116, 201904978 (2019).

14. Boilard, E. Extracellular vesicles and their content in bioactive lipid mediators: more than a sack of microRNA. J Lipid Res 59, 2037–2046 (2018).

15. Boudreau, L. H. et al. Platelets release mitochondria serving as substrate for bactericidal group IIA-secreted phospholipase A2 to promote inflammation. Blood 124, (2014).

16. Marcoux, G. et al. Platelet-derived extracellular vesicles convey mitochondrial DAMPs in platelet concentrates and their levels are associated with adverse reactions. Transfusion 59, 2403–2414 (2019).

17. Pelletier, M. et al. Platelet extracellular vesicles and their mitochondrial content improve the mitochondrial bioenergetics of cellular immune recipients. Transfusion (Paris) 63, 1983–1996 (2023).

18. Pelletier, M. et al. Platelet extracellular vesicles and their mitochondrial content improve the mitochondrial bioenergetics of cellular immune recipients. Transfusion (Paris) 63, 1983–1996 (2023).

19. Puhm, F. et al. Mitochondria Are a Subset of Extracellular Vesicles Released by Activated Monocytes and Induce Type I IFN and TNF Responses in Endothelial Cells. Circ Res 125, 43–52 (2019).

20. Melchinger, H., Jain, K., Tyagi, T. & Hwa, J. Role of Platelet Mitochondria: Life in a Nucleus-Free Zone. Front Cardiovasc Med 6, 1–11 (2019).

21. Vakifahmetoglu-Norberg, H., Ouchida, A. T. & Norberg, E. The role of mitochondria in metabolism and cell death. Biochemical and Biophysical Research Communications vol. 482 426–431 (2017).

22. Jin, P. et al. Platelets Facilitate Wound Healing by Mitochondrial Transfer and Reducing Oxidative Stress in Endothelial Cells. Oxid Med Cell Longev 2023, (2023).

23. Levoux, J. et al. Platelets Facilitate the Wound-Healing Capability of Mesenchymal Stem Cells by Mitochondrial Transfer and Metabolic Reprogramming. Cell Metab 33, 283-299.e9 (2021).

24. Torralba, D., Baixauli, F. & Sánchez-Madrid, F. Mitochondria Know No Boundaries: Mechanisms and Functions of Intercellular Mitochondrial Transfer. Front Cell Dev Biol 4, 107 (2016).

25. Chen, J., Zhong, J., Wang, L. & Chen, Y. Mitochondrial Transfer in Cardiovascular Disease: From Mechanisms to Therapeutic Implications. Front Cardiovasc Med 8, 1–15 (2021).

26. Qin, Y. et al. The Functions, Methods, and Mobility of Mitochondrial Transfer Between Cells. Frontiers in Oncology vol. 11 (2021).

27. Veilleux, V., Pichaud, N., Boudreau, L. H. & Robichaud, G. A. Mitochondria transfer by platelet-derived microparticles regulates breast cancer bioenergetic states and malignant features. Molecular Cancer Research (2023)

28. Chen, L. et al. Astrocyte mitochondria: Potential therapeutic targets for epilepsy. Heliyon vol. (2024).

29. Iorio, R., Petricca, S., Mattei, V. & Delle Monache, S. Horizontal mitochondrial transfer as a novel bioenergetic tool for mesenchymal stromal/stem cells: molecular mechanisms and therapeutic potential in a variety of diseases. Journal of Translational Medicine vol. 22 (2024).

30. Zhou, Z. et al. Extracellular vesicles activated cancer-associated fibroblasts promote lung cancer metastasis through mitophagy and mtDNA transfer. Journal of Experimental and Clinical Cancer Research 43, (2024).

31. Baharvand, F. et al. Safety and efficacy of platelet-derived mitochondrial transplantation in ischaemic heart disease. Int J Cardiol 410, (2024).

32. Caicedo, A. et al. The diversity and coexistence of extracellular mitochondria in circulation: A friend or foe of the immune system. Mitochondrion vol. 58 270–284 (2021).

33. Cappelletto, A. et al. SARS-CoV-2 Spike protein activates TMEM16F-mediated platelet procoagulant activity. Front Cardiovasc Med 9, 1–16 (2023).

34. Allan, H. E. et al. Proteome and functional decline as platelets age in the circulation. Journal of Thrombosis and Haemostasis 00, 1–18 (2021).

35. Ferreira, P. M. et al. Mode of induction of platelet-derived extracellular vesicles is a critical determinant of their phenotype and function. Sci Rep 10, (2020).

36. Headland, S. E., Jones, H. R., D’Sa, A. S. V., Perretti, M. & Norling, L. V. Cutting-edge analysis of extracellular microparticles using imagestreamx imaging flow cytometry. Sci Rep 4, (2014).

37. Jeon, J. H., Hong, C. W., Kim, E. Y. & Lee, J. M. Current understanding on the metabolism of neutrophils. Immune Network vol. 20 1–13 (2020).

38. Bao, Y. et al. Mitochondria regulate Neutrophil activation by generating ATP for Autocrine Purinergic signaling. Journal of Biological Chemistry 289, 26794–26803 (2014).

39. Miralda, I., Uriarte, S. M. & McLeish, K. R. Multiple Phenotypic Changes Define Neutrophil Priming. Front Cell Infect Microbiol 7, 217 (2017).

40. Abdel-Salam, B. K. A. & Ebaid, H. Expression of CD11b and CD18 on polymorphonuclear neutrophils stimulated with interleukin-2. Cent Eur J Immunol 39, 209–15 (2014).

41. Lakschevitz, F. S. et al. Identification of neutrophil surface marker changes in health and inflammation using high-throughput screening flow cytometry. Exp Cell Res 342, 200–209 (2016).

42. Hann, J., Bueb, J. L., Tolle, F. & Bréchard, S. Calcium signaling and regulation of neutrophil functions: Still a long way to go. Journal of Leukocyte Biology vol. 107 285–297 (2020).

43. Papayannopoulos, V. Neutrophil extracellular traps in immunity and disease. Nature Reviews Immunology vol. 18 134–147 (2018).

44. Hidalgo, A. et al. Neutrophil extracellular traps: from physiology to pathology. Cardiovascular Research vol. 118 2737–2753 (2022).

45. Zhang, Q. et al. Circulating mitochondrial DAMPs cause inflammatory responses to injury. Nature 464, 104–107 (2010).

46. Wang, Y. et al. Histone hypercitrullination mediates chromatin decondensation and neutrophil extracellular trap formation. Journal of Cell Biology 184, 205–213 (2009).

47. Sorvillo, N., Cherpokova, D., Martinod, K. & Wagner, D. D. Extracellular DNA net-works with dire consequences for health. Circulation Research vol. 125 470–488 (2019).

48. Kirchner, T. et al. The impact of various reactive oxygen species on the formation of neutrophil extracellular traps. Mediators Inflamm 2012, (2012).

49. Azzouz, D., Khan, M. A. & Palaniyar, N. ROS induces NETosis by oxidizing DNA and initiating DNA repair. Cell Death Discov 7, (2021).

50. Doroshenko, T. et al. Phagocytosis-induced modulation of human neutrophil chemotaxis receptors. Blood 58, 228–236 (2002).

51. Doroshenko, T. et al. Phagocytosing Neutrophils Down-Regulate the Expression of Chemokine Receptors CXCR1 and CXCR2. Blood. 58(2): 228–236 (2002).

52. Manfredi, A. A., Ramirez, G. A., Rovere-Querini, P. & Maugeri, N. The neutrophil’s choice: Phagocytose vs make neutrophil extracellular traps. Frontiers in Immunology, 9(288):1–13 (2018).

53. Gonzalez-Delgado, R. et al. Role of circulating mitochondria in venous thrombosis in glioblastoma. Journal of Thrombosis and Haemostasis 21, 2202–2212 (2023).

54. Vijayan, V. et al. Extracellular release of damaged mitochondria induced by prehematopoietic stem cell transplant conditioning exacerbates GVHD. Blood Adv 8, 3691–3704 (2024).

55. Michailidou, D., Giaglis, S. & Dale, G. L. The platelet-mitochondria nexus in autoimmune and musculoskeletal diseases. Clinical Immunology vol. 267 (2024).

56. Urban, C. F., Reichard, U., Brinkmann, V. & Zychlinsky, A. Neutrophil extracellular traps capture and kill Candida albicans and hyphal forms. Cell Microbiol 8, 668–676 (2006).

57. Meier, A., Sakoulas, G., Nizet, V. & Ulloa, E. R. Neutrophil Extracellular Traps: An Emerging Therapeutic Target to Improve Infectious Disease Outcomes. Journal of Infectious Diseases, 230: 514–521 (2024).

58. Martinod, K. & Wagner, D. D. Thrombosis: tangled up in NETs. (2014) doi:10.1182/blood.

59. Gidlöf, O. et al. Platelets activated during myocardial infarction release functional miRNA, which can be taken up by endothelial cells and regulate ICAM1 expression. Blood 121, 3908– 17, S1-26 (2013).

