## Supplemental Figures for "Platelet mitochondrial transfer via extracellular vesicles modulates neutrophil phenotype and function"

### Supplemental Data

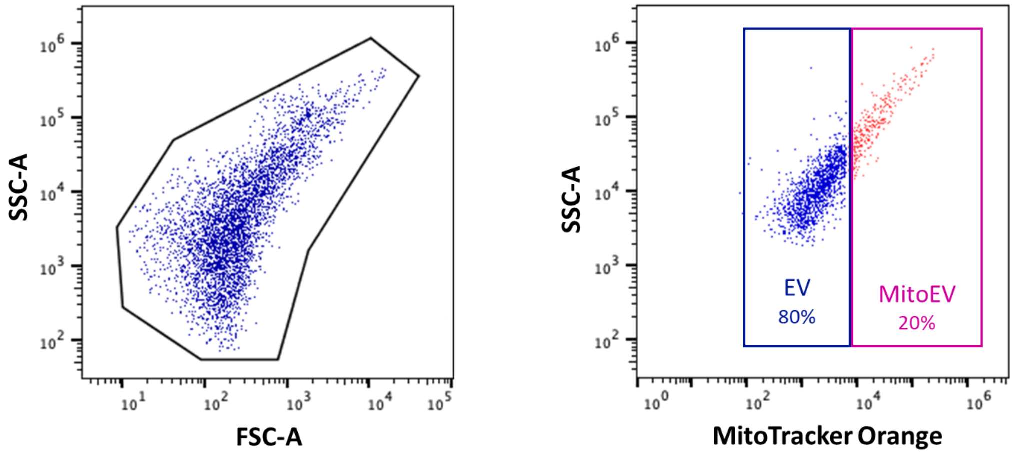


**Supplemental Figure 1: Representative Gating Strategy for Sorting of Platelet Extracellular Vesicles.** Platelet extracellular vesicles were sorted into two subpopulations based on the fluorescence of MitoTracker Orange; those without a mitochondrion (EV, 80%; blue) and those with a mitochondrion (mitoEV, 20%; magenta).

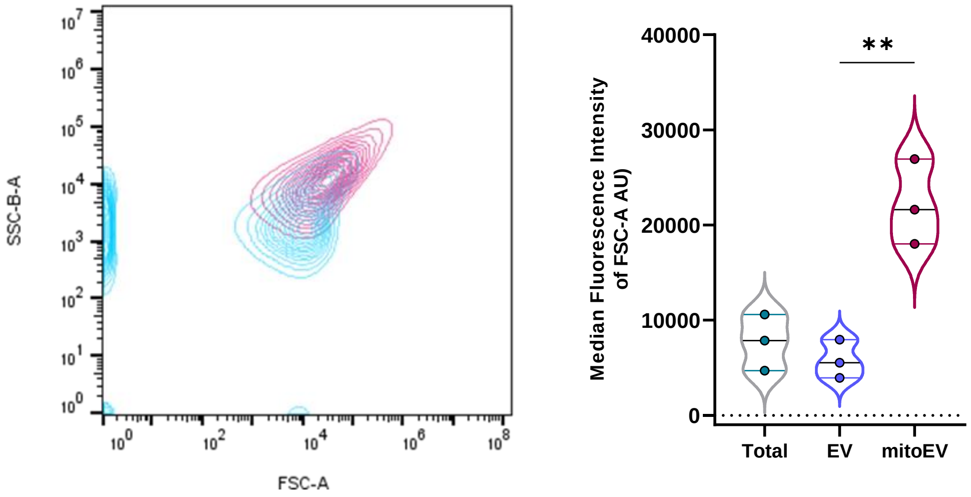


**Supplemental Figure 2: FSC/SSC of Extracellular Vesicles. (A)** Representative flow plot showing Forward Scatter vs. Side Scatter as an indicator of extracellular vesicle size. **(B)** Quantification of forward scatter as an indicator of platelet extracellular vesicle size split into those containing a mitochondria (mitoEVs; magenta) and platelet extracellular vesicles without mitochondria (EVs; blue) analysed using the Cytek Aurora.

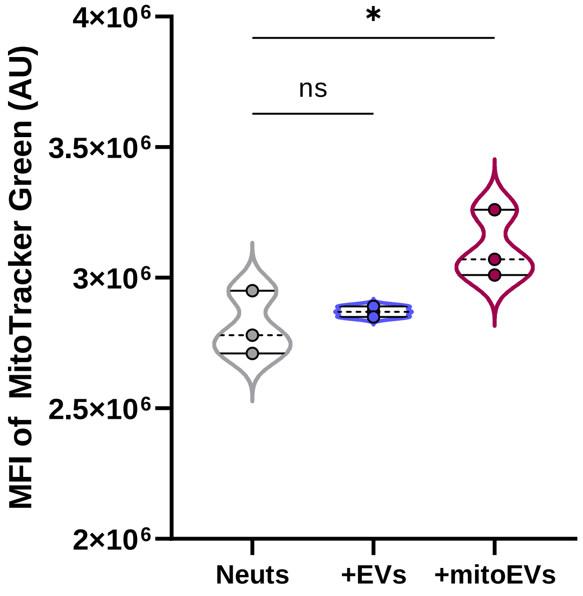


**Supplemental Figure 3: Analysis of neutrophil mitochondrial content.** Flow cytometric analysis of neutrophil mitochondrial mass (MitoTracker Green fluorescence) either alone (grey) or following incubation with EVs (blue) or mitoEV (magenta; n=3, $p<0.05$)

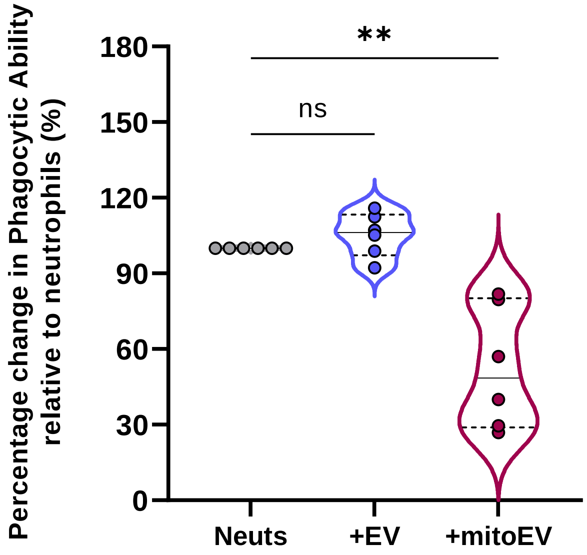


**Supplemental Figure 4: Characterisation of phagocytosis by neutrophils following incubation with platelet extracellular vesicles.** Flow cytometric analysis of neutrophil phagocytosis of pHrodo$^{TM}$ *E.coli b*ioparticles following incubation with PMVs (blue) or mitoPMVs (magenta) compared to neutrophils alone (n=6, $p<0.05$)
